## Supplemental Materials for "Voxel-Matching NORDIC: Non-local patch formation by time-series similarity increases tSNR in high-resolution BOLD fMRI"

### SUPPLEMENTARY MATERIALS

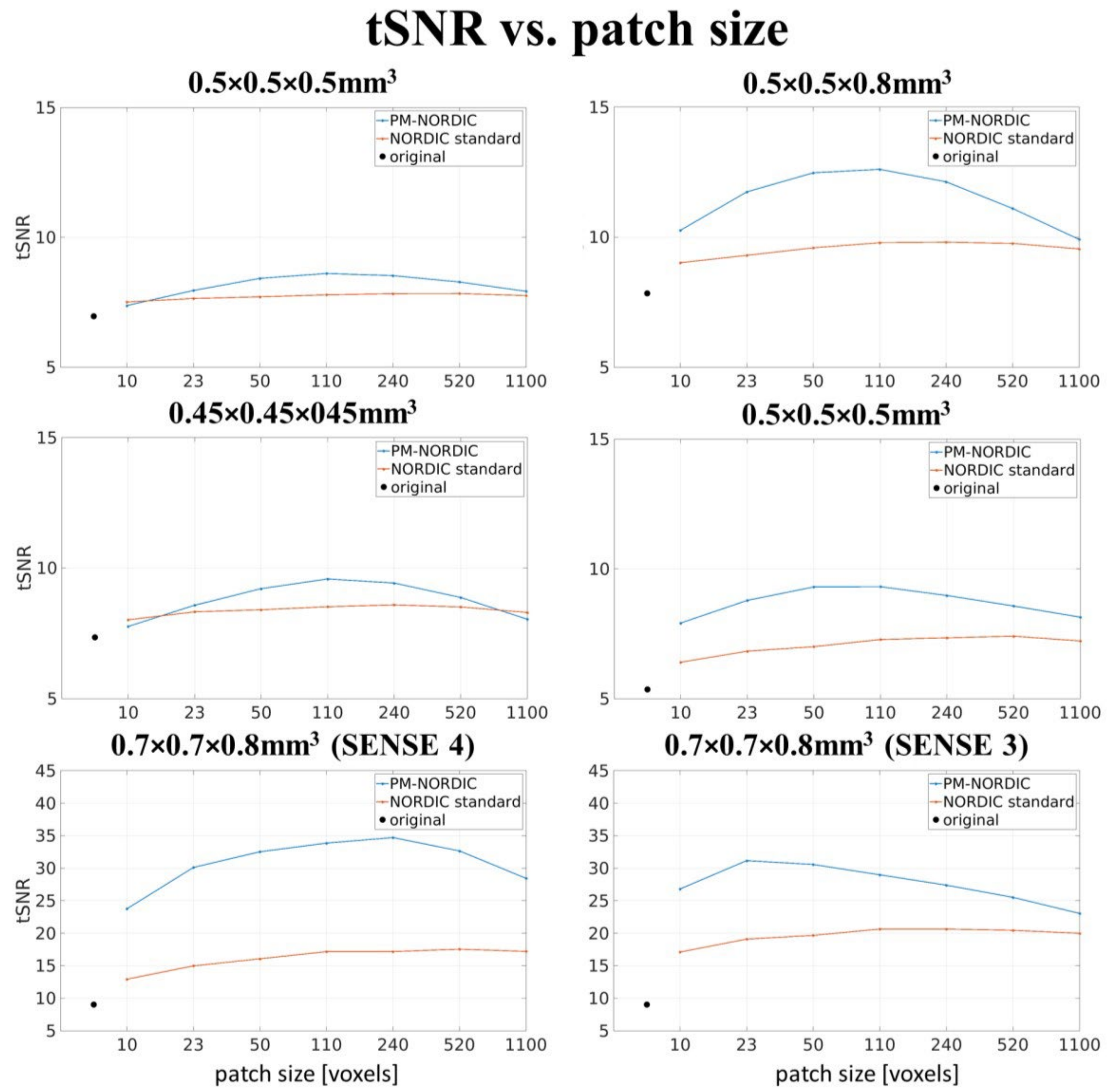


**Supplementary Figure 1.** Mean tSNR as a function of patch size for Standard-NORDIC and VM-NORDIC. The denoising performance of VM-NORDIC is more sensitive to the chosen patch size. Also, The curves for VM-NORDIC show clear maxima representing the optimal patch sizes.

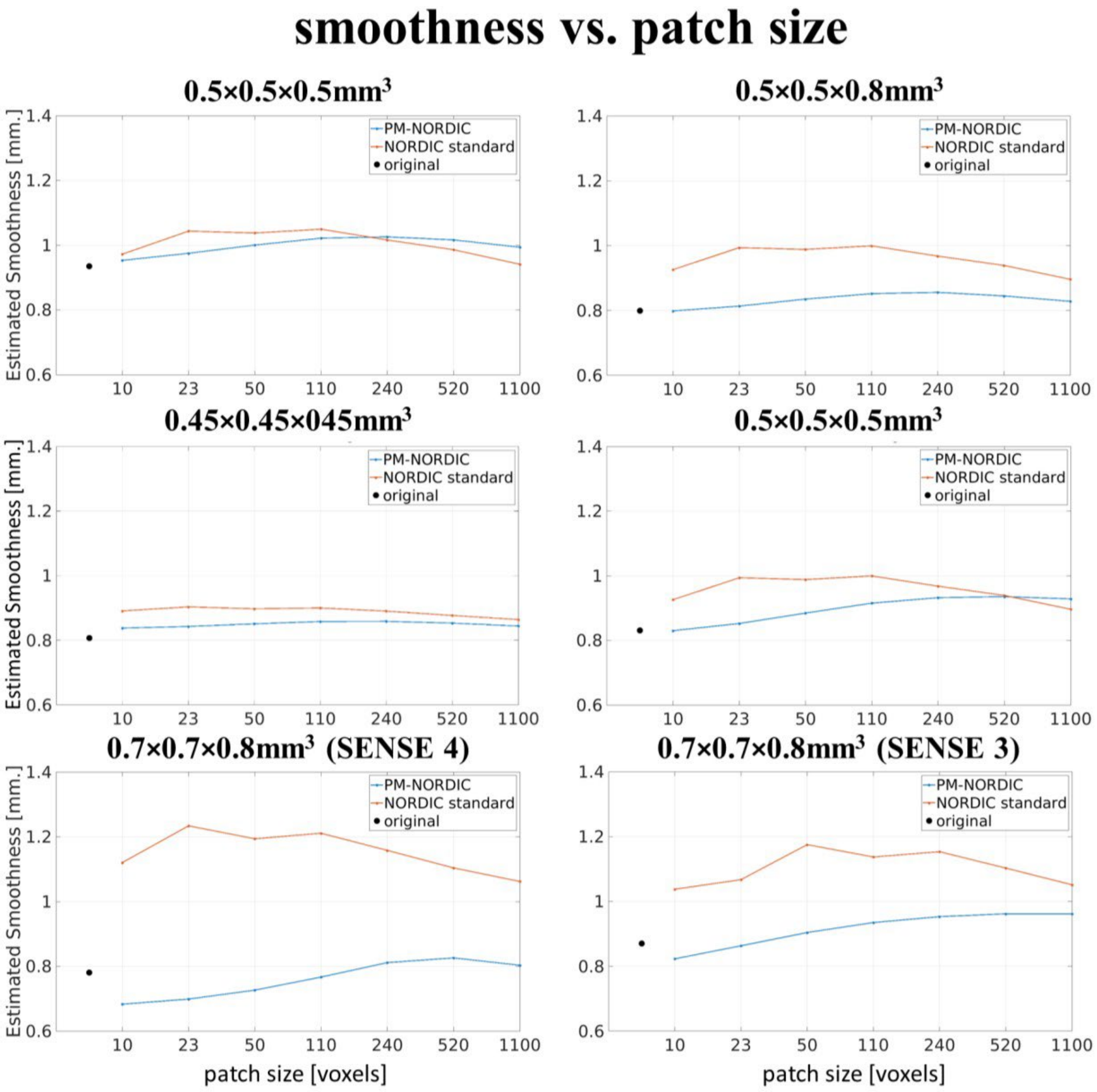


**Supplementary Figure 2.** Global smoothness estimates (FWHM) as a function of patch size for Standard-NORDIC and VM-NORDIC. VM-NORDIC does not significantly increase spatial smoothing and, for certain patch sizes, even leads to smoothness estimates lower than the originals by counteracting smoothing signal leakage.

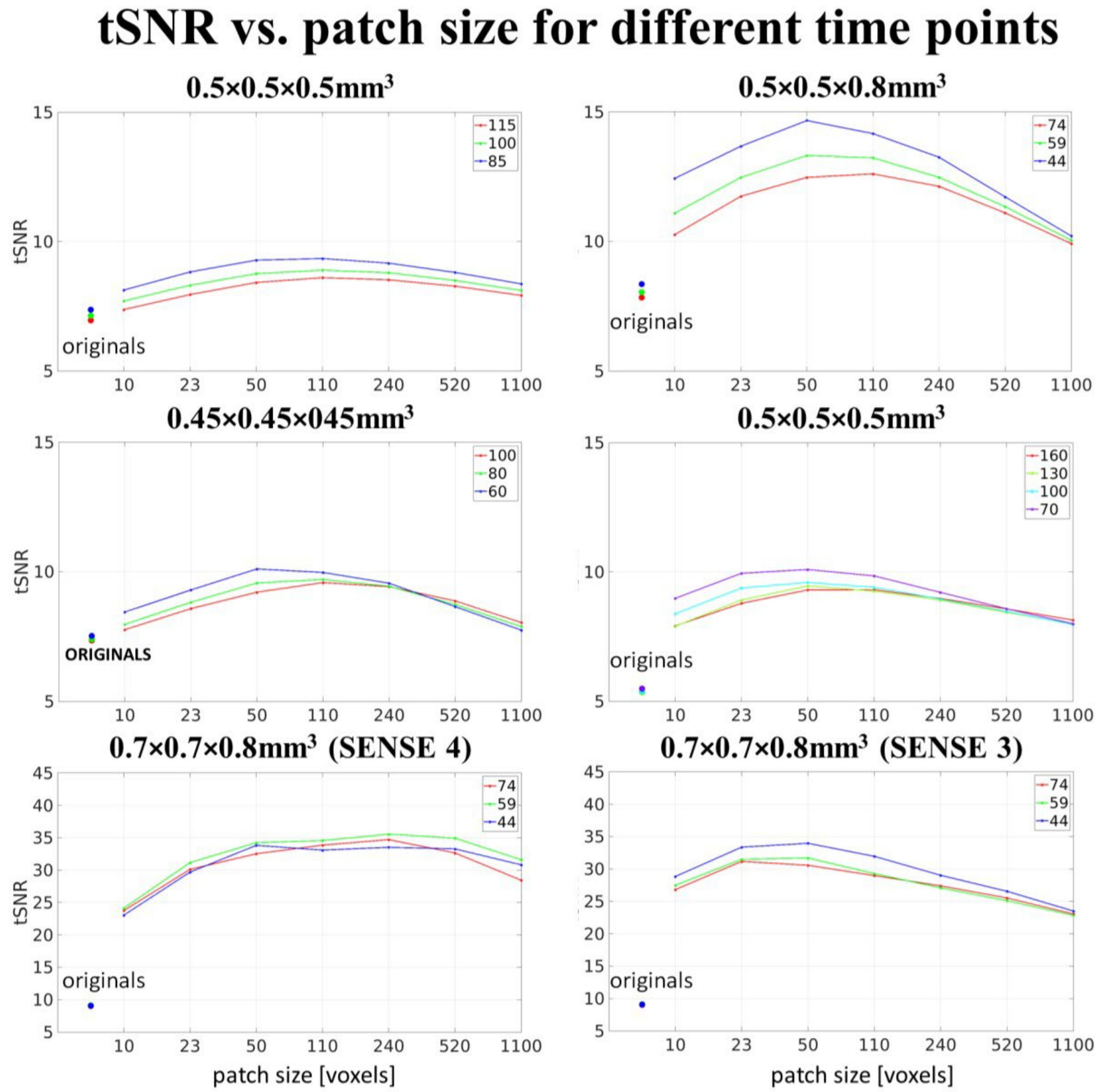


**Supplementary Figure 3.** Mean tSNR as a function of patch size for VM-NORDIC denoised datasets with different numbers of time points. The optimal patch size slightly increases with increasing time points. Nonetheless, the shift is not significant as long as the number of time points is within a regular range. Additionally, shorter datasets show the largest tSNR gains.

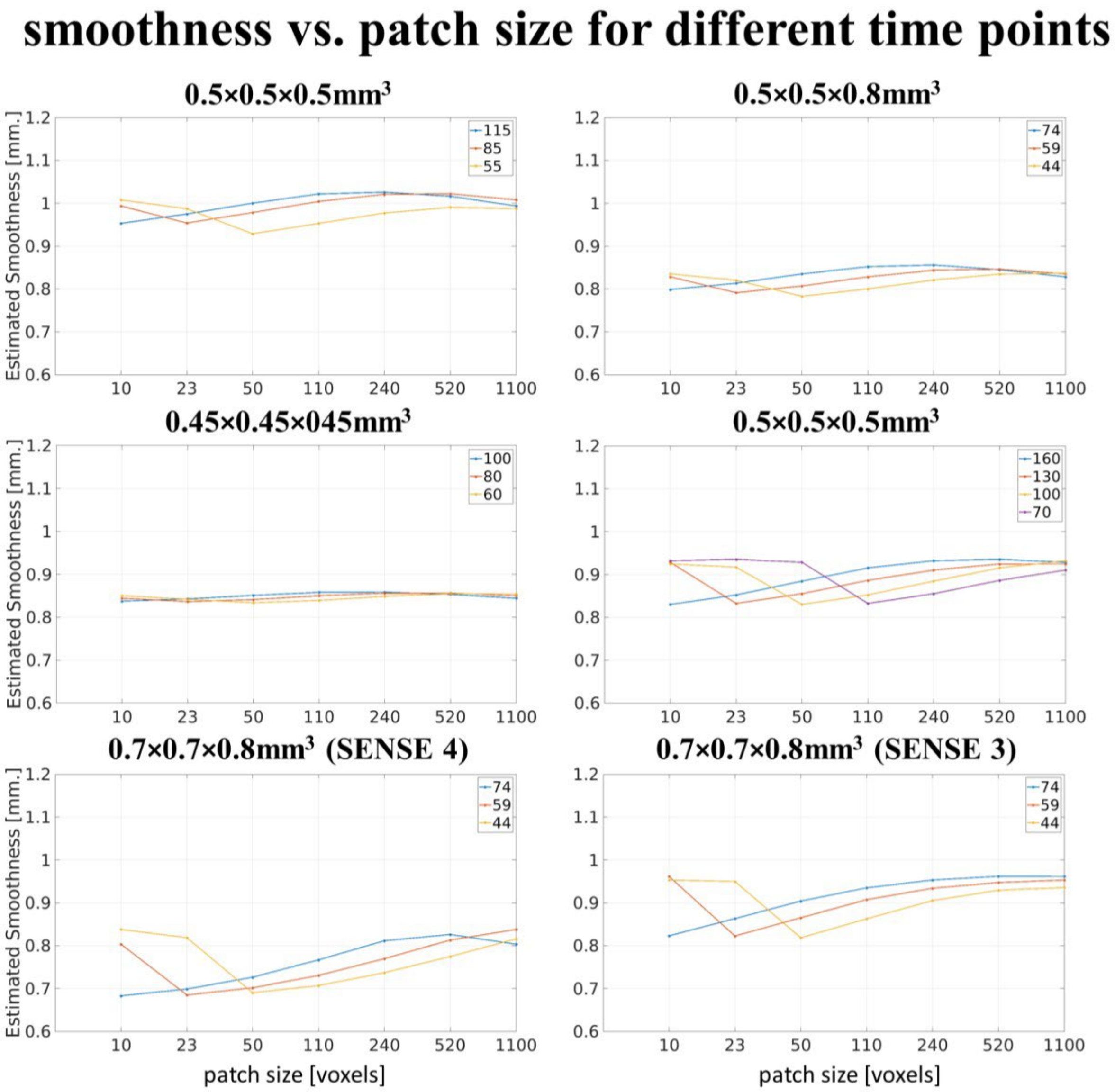


**Supplementary Figure 4.** Global smoothness estimates (FWHM) as a function of patch size for VM-NORDIC denoised datasets with different numbers of time points. The degree of spatial blurring exhibits small but irrelevant fluctuations as the number of time points of the dataset changes.

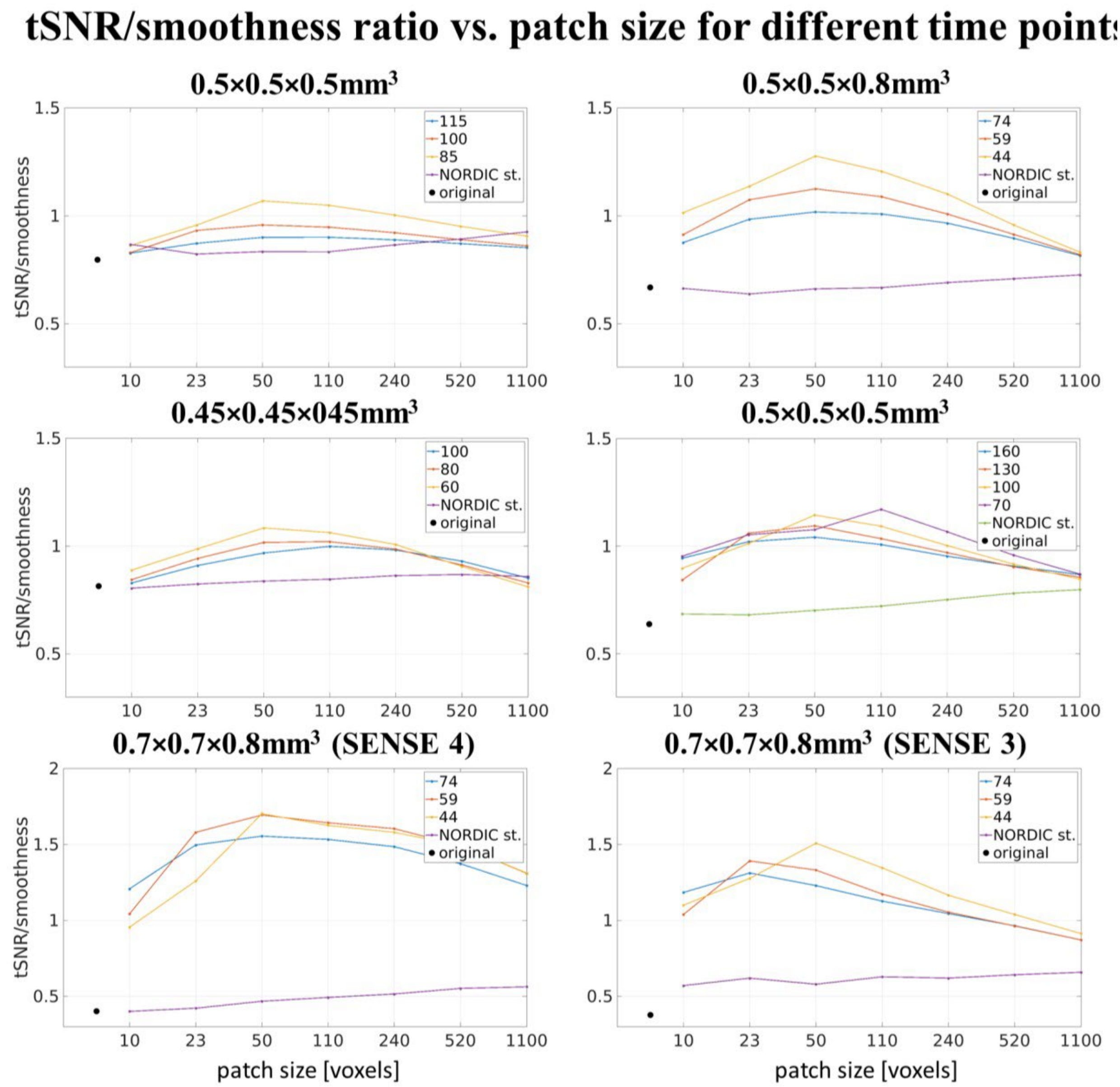


**Supplementary Figure 5.** Normalized tSNR over normalized global smoothness estimate (FWHM) ratio as a function of patch size for VM-NORDIC denoised datasets with different number of time points. This metric shows which patch size guarantees the best denoising performance in terms of the trade-off between nosie removal and induced spatial smoothness. The curves indicate that VM-NORDIC has a clear advantage over Standard-NORDIC, and that the optimal patch sizes range from 20 up to 240 timeseries per patch.
